# Nanosecond laser-driven proton FLASH spares normal tissue cells by sustaining mitochondrial homeostasis and attenuating ferroptosis

**DOI:** 10.64898/2026.03.13.711531

**Authors:** Changsheng Shao, Yu Zhang, Pengzhi He, Xin Yu, Wenjie Peng, Junfeng Chen, Haotian Hu, Yiying Wang, Mei Xiao, Chao Liu, Li Sui, Tianyuan Dai, Xiangkui Mu, Xianghong Jia, Jianhui Bin, Qing Huang

## Abstract

Radiotherapy’s clinical utility remains fundamentally constrained by the collateral damage to healthy tissues. Ultra-high-dose-rate (UHDR) irradiation, or, FLASH-radiotherapy (FLASH-RT) has emerged as a transformative paradigm to mitigate such toxicity. However, the biological effects of FLASH-RT on the high-efficiency of tumor killing and normal tissue sparing remain poorly understood. In this work, we utilized a petawatt-class laser-plasma acceleration (LPA) platform to deliver discrete 12.9-nanosecond proton pulses at an extreme instantaneous dose rate of 1.94×10^7^ Gy/s. This temporal singularity achieved a profound sparing effect in normal bronchial epithelial cells, evidenced by a nine-fold reduction in the lethal α coefficient (from 0.47 to 0.05 Gy^-1^), while maintaining full tumoricidal potency against lung adenocarcinoma. Mechanistically, we demonstrated that LPA-FLASH could effectively bypass the ATF3-mediated stress response and circumvent the subsequent ferroptotic cascade. This molecular evasion could preserve the mitochondrial cristae integrity and trigger an adaptive bioenergetic ATP surge—a hallmark of metabolic resilience exclusively in healthy tissue cells. Therefore, our findings identify ferroptosis-mediated mitochondrial integrity as a unifying framework for selective normal-tissue protection at the physical limits of radiation delivery, and establish LPA-FLASH-RT as a potent, compact modality for next-generation oncology.

---

Radiotherapy becomes a cornerstone of comprehensive cancer treatment; however, its clinical efficacy is limited by a narrow therapeutic window resulting from collateral damage to healthy stromal tissues [**1**]. FLASH radiotherapy (FLASH-RT) is a technique that administers ultra-high-dose-rate (UHDR) irradiation in less time without compromising tumor treatment [**2–4**]. Preclinical studies across diverse animal models have consistently demonstrated that FLASH-RT significantly mitigates acute and late complications in normal tissues while maintaining tumor control comparable to conventional radiotherapy (CONV-RT), a paradigm-shifting biological selectivity termed the “FLASH effect” [**5–7**].

Despite the promising therapeutic potential of FLASH-RT, the underlying radiobiological mechanisms remain a subject of intense debate. Current hypotheses encompass rapid oxygen depletion leading to transient radiochemical hypoxia [**8**], alteration of radical-radical recombination [**9**], and the preservation of mitochondrial function [**10**]. Recent evidence also suggests that ferroptosis—an iron-dependent form of regulated cell death characterized by lipid peroxidation **[11**]—may be a pivotal executioner of the FLASH-mediated sparing effect [**12,13**].

A defining characteristic of FLASH-RT is its extreme instantaneous dose rate. To date, most FLASH research has utilized traditional radiofrequency linear accelerators or cyclotrons operating in the microsecond regime. Importantly, recent advances in compact laser-plasma acceleration (LPA) have opened a fundamentally different physical regime. Laser-driven proton sources utilize petawatt-class femtosecond laser pulses to generate proton beams with durations in the picosecond-to-nanosecond range, achieving instantaneous dose rates exceeding 10^9^ Gy/s—six orders of magnitude higher than conventional FLASH systems [**14–15**].

In this study, by employing a high-repetition-rate laser-driven proton platform at the Shanghai Institute of Optics and Fine Mechanics (SIOM) (so-called SIOM HLDP platform). we attempted to reveal the LPA-FLASH effect in the petawatt–picosecond regime and explore the molecular underpinnings. By delivering protons in discrete 12.9 ns pulses with extreme instantaneous dose rates, we systematically compared the biological impact on human A549 lung adenocarcinoma cells and BEAS-2B normal bronchial epithelial cells against conventional (CONV) clinical protons. Through an integrated approach—incorporating 3D light-sheet fluorescence microscopy for subcellular morphometric quantification, transmission electron microscopy (TEM) for ultrastructural analysis, and functional mitochondrial assays—we demonstrated that LPA-FLASH-RT could give rise to a differential biological response characterized by the selective preservation of mitochondrial cristae in normal cells. By characterizing the spatio-temporal dynamics of the special ATF3-mediated ferroptotic axis, we revealed that the temporal singularity of laser-driven protons could probe a transcriptional blind spot, effectively bypassing lethal oxidative stress cascades. Therefore, this study may help to establish LPA-FLASH-RT as a potent modality that shifts the focus of radiobiology from DNA-centric damage to a framework of metabolic resilience, and providing a mechanistically defined roadmap for the next-generation radiotherapy.

## Results

### LPA-FLASH-RT establishe a superior therapeutic window by selectively sparing normal-tissue

To investigate whether the extreme temporal compression of laser-driven protons translates into a biological advantage, we utilized the 200 TW laser-driven proton source at the SIOM HLDP platform. This system generates protons in discrete, ultra-short pulses (Δ*t* = 12.9 ns), delivering 0.25 Gy per shot and achieving an extreme instantaneous dose rate (IDR) of 1.94×10^7^ Gy/s (hereafter referred to as LPA-FLASH-RT). We directly compared this regime with a conventional proton accelerator operating at a standard clinical dose rate of 0.08 Gy/s (hereinafter CONV-RT) (**Fig. 1a, b**).

**Figure 1.**
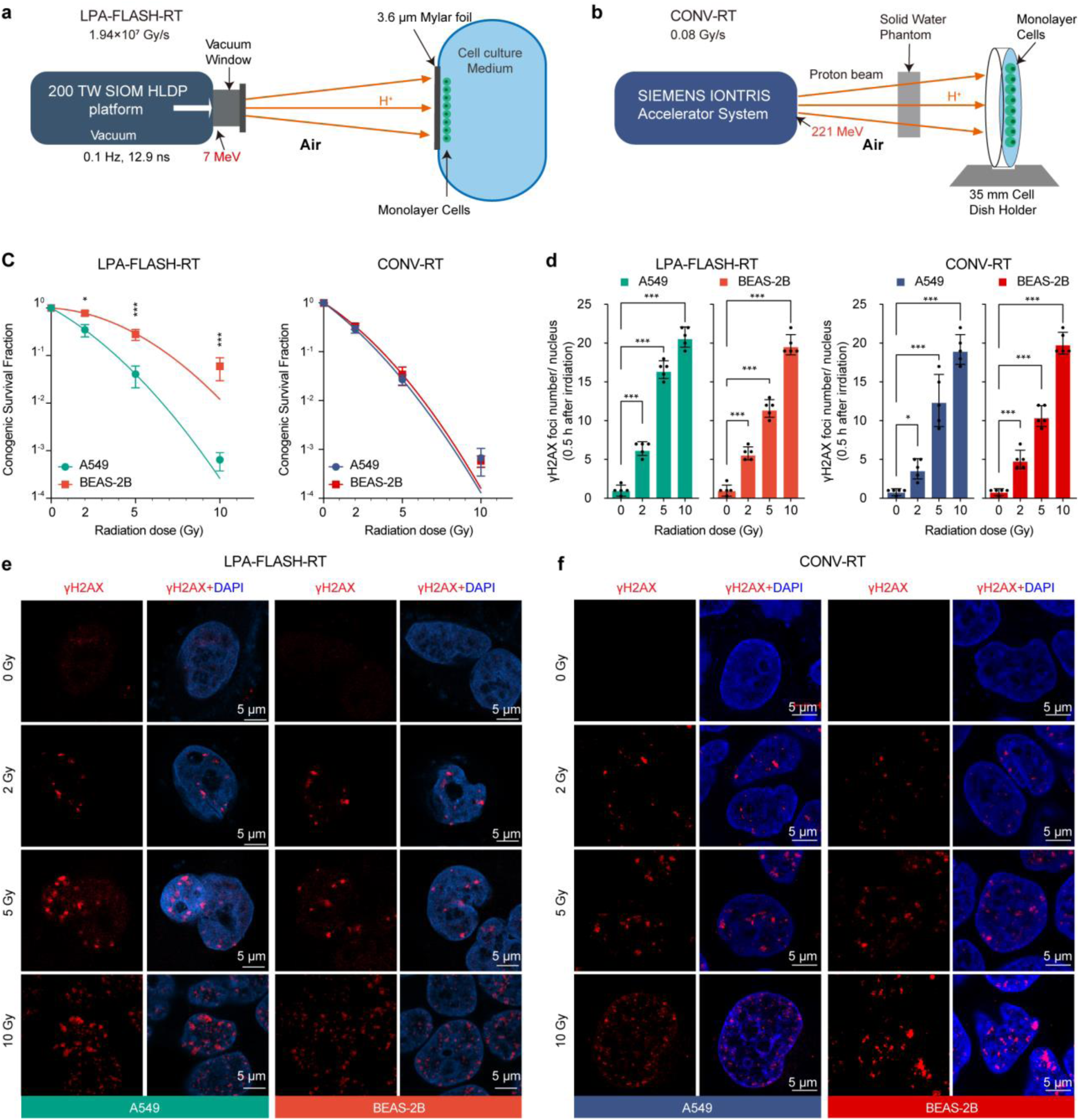
Differential biological effects of laser-driven FLASH versus conventional proton irradiation. **(a, b)** Schematic illustrations of the experimental setups for the laser-driven FLASH proton irradiation (LPA-FLASH-RT) system **(a)**and the conventional proton irradiation (CONV-RT) system**(b)**. (**c**) Clonogenic survival fractions (SF) of BEAS-2B normal lung epithelial cells and A549 lung adenocarcinoma cells. The left panel shows survival curves under LPA-FLASH-RT, and the right panel shows survival curves under CONV-RT. Data were fitted using the linear-quadratic model (n=3 independent experiments). (**d)** Quantification of DNA double-strand breaks as measured by the number of γ-H2AX foci per cell (n=5, random fields). **(e, f)** Representative immunofluorescence images of γ-H2AX foci (red) and DAPI-stained nuclei (blue) in cells at 0.5 h post-irradiation with LPA-FLASH-RT **(e)** and CONV-RT **(f)**. Scale bar, 5 μm. Data are presented as mean ± s.d. Statistical significance was determined using one-way ANOVA followed by Tukey’s multiple comparisons test; ns, no significant difference (p > 0.05); *, significant difference (p < 0.05); **, very significant difference (p < 0.01); ***, highly significant difference (p < 0.001).

Clonogenic survival assays on human A549 lung cancer cells and BEAS-2B normal epithelial cells revealed that LPA-FLASH-RT creates a profound therapeutic window by selectively sparing healthy tissue while maintaining full malignant-cell kill. In A549 tumor cells, the survival curves and linear-quadratic (LQ) parameters were nearly indistinguishable between the two modalities (**Fig. 1c**). The lethal α value for LPA-FLASH-RT (0.46 Gy⁻¹) remained comparable to that of CONV-RT (0.55 Gy⁻¹, p>0.05), confirming that the nine-order-of-magnitude increase in dose rate does not compromise tumoricidal potency.

In sharp contrast, BEAS-2B normal cells exhibited a dramatic survival advantage under LPA-FLASH-RT. The α value, representing non-repairable lethal damage, plummeted nine-fold from 0.47 Gy⁻¹ under CONV-RT to 0.052 Gy⁻¹ under LPA-FLASH-RT (p < 0.001; **Fig. 1c**), while the α/β ratio collapsed from 11.87 Gy to 1.35 Gy. These data confirm that the extreme temporal compression of laser-protons yields potent normal-tissue sparing without compromising tumoricidal efficacy.

### LPA-FLASH-RT irradiation mitigates initial genomic stress in normal cells

The profound nine-fold reduction in the lethal α value of normal cells suggested that the nanosecond pulse might circumvent the induction of complex genomic lesions. To test this biological decoupling hypothesis at the earliest stage of injury, we quantified γ-H2AX foci formation 0.5 h post-irradiation, specifically focusing on the 5 Gy dose point where the divergence between normal and malignant cell survival was most pronounced. In A549 tumor cells, both LPA-FLASH-RT and CONV-RT induced extensive and dense foci distribution, with 10 Gy irradiation increasing the mean foci count to 20.8 ± 1.2 and 19.2 ± 1.9 per nucleus, respectively (p > 0.05).

For normal BEAS-2B cells, while CONV-RT produced a high density of DNA damage (20.1 ± 1.4 foci at 10 Gy), the FLASH-irradiated group showed a significantly lower rate of foci accumulation at lower clinical doses. At 2 Gy, LPA-FLASH-RT induced only 6.4 ± 0.9 foci in A549 cells, whereas BEAS-2B cells under identical FLASH conditions showed a marked trend toward more efficiently managed genomic stress (p < 0.001 relative to tumor cells; **Fig. 1d-e**).

To further delineate the dose-response relationship of genomic stress, we focused on the 5 Gy dose point—a clinically relevant threshold for normal tissue tolerance. At 5 Gy, LPA-FLASH-RT elicited a strikingly divergent response between cell types: while A549 tumor cells sustained extensive DNA double-strand breaks (DSBs) comparable to conventional levels, BEAS-2B normal cells exhibited a significant reduction in γ-H2AX foci formation (p < 0.001 relative to A549). Quantitatively, the initial lesion density in FLASH-irradiated normal cells at 5 Gy was significantly lower than that of their tumor counterparts, suggesting that the ultra-high-dose-rate pulse triggers a rapid genomic sparing effect that is uniquely operative in the normal tissue background ( **Fig. 1d**). This selective safeguarding of genomic integrity in normal stroma aligns with the enhanced survival observed in the clonogenic assays, suggesting a divergent early stress-sensing mechanism under UHDR kinetics.

### LPA-FLASH-RT bypass the pro-ferroptotic ATF3 stress-response axis in normal epithelial cells

To bridge the gap between cell survival and genomic integrity, we investigated the biochemical stress dynamics following irradiation, beginning with the immediate production of reactive oxygen species (ROS). At one-hour post-irradiation, marking the onset of the post-irradiation biochemical window, total cellular ROS levels were measured to evaluate the initial chemical impact on the cells. In A549 tumor cells, both radiation types caused a large increase in ROS that depended on the dose. Quantitatively, at 10 Gy, LPA-FLASH-RT increased ROS intensity to 908 ± 9.3 RFU and CONV-RT increased it to 1012 ± 103 RFU (p < 0.001 for both). However, we saw a different result in normal BEAS-2B cells (**Fig. 2a**). While CONV-RT caused ROS in normal cells to rise significantly to 895 ± 13 RFU at 10 Gy (p < 0.001), LPA-FLASH-RT kept ROS levels low at 527± 12 RFU, which is near the baseline.

**Figure 2.**
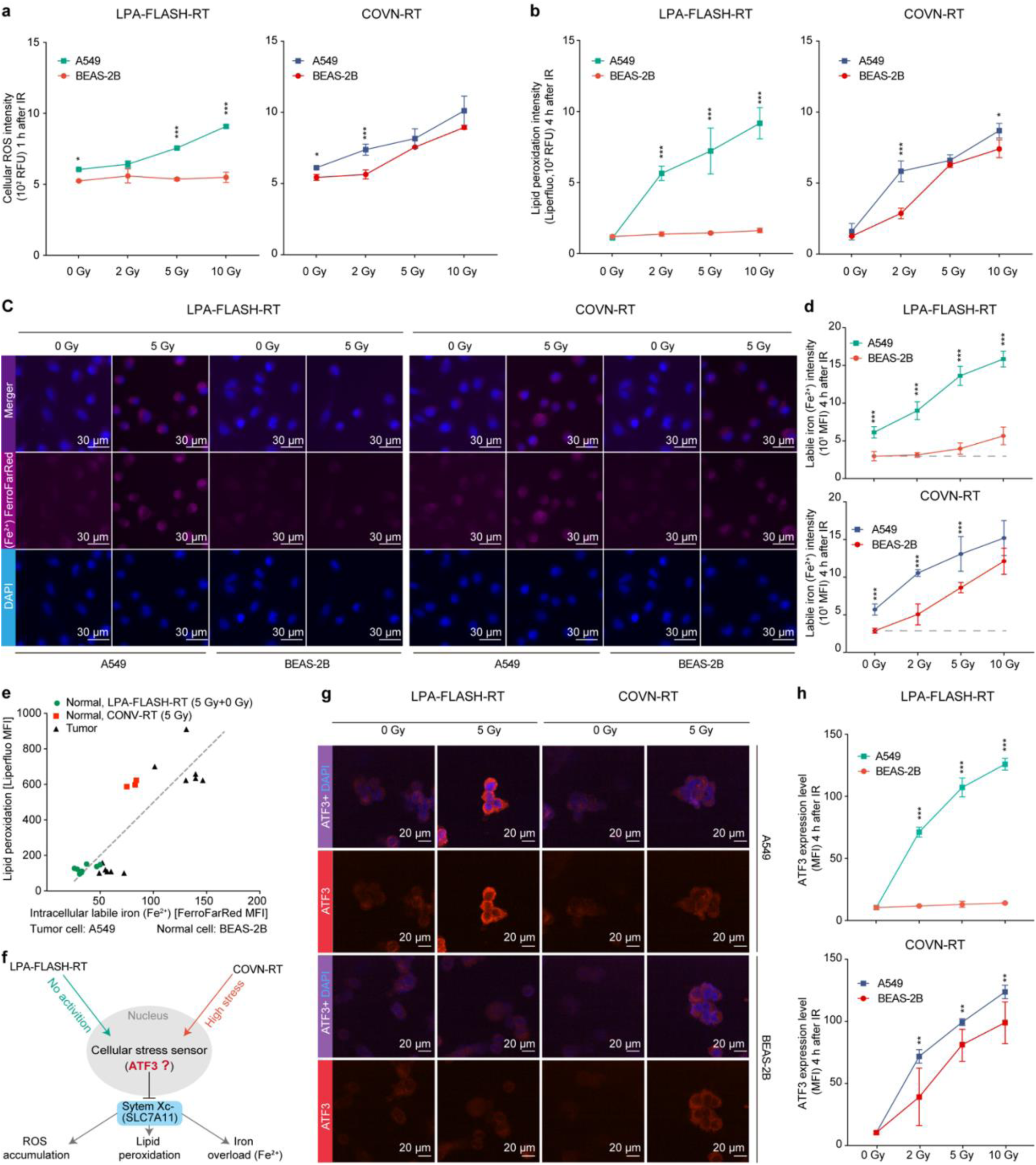
FLASH irradiation mitigates ferroptosis-related oxidative stress and iron overload by bypassing ATF3 activation. **(a, b)** Quantification of intracellular oxidative stress markers in BEAS-2B and A549 cells. Intracellular ROS levels **(a)** were assessed using the H_2_DCFDA probe, and lipid peroxidation **(b)** was measured using the Liperfluo probe. In both **(a)** and **(b)**, the left panels show results for LPA-FLASH-RT and the right panels for CONV-RT. Data are expressed as relative fluorescence units. **(c)** Representative fluorescence microscopy images of the labile iron pool (Fe^2+^) levels at 4 h post-irradiation. **(d)** The upper panel displays data for FLASH-RT, and the lower panel for CONV-RT. Scale bar, 30 μm. **(e)** Pearson correlation analysis between intracellular Fe^2+^levels and lipid peroxidation (Liperfluo intensity). **(f)** Schematic hypothesis illustrating ATF3 as a potential cellular stress sensor that triggers the cascade of ROS accumulation, lipid peroxidation, and iron overload. **(g)** Representative immunofluorescence images showing ATF3 expression and localization. **(h)** Dose-dependent quantification of ATF3 expression levels at 4 h post-irradiation (0, 2, 5, and 10 Gy). The upper panel corresponds to LPA-FLASH-RT, and the lower panel to CONV-RT. Data are presented as mean ± s.d. derived from 3 independent experiments. Statistical significance was analyzed using one-way ANOVA; ns, no significant difference (p > 0.05); *, significant difference (p < 0.05); **, very significant difference (p < 0.01); ***, highly significant difference (p < 0.001).

We reasoned that this initial attenuation of oxidative stress under LPA-FLASH-RT might prevent the cell from crossing a critical signaling threshold during the subsequent 1–4 hour biochemical decision window. To test whether this early protection prevents the execution of ferroptotic cell death, we measured lipid peroxidation (LPO) at 4 hours after radiation using the Liperfluo probe. In A549 cancer cells, both LPA-FLASH-RT and CONV-RT induced heavy lipid damage. Intensity levels reached 918 ± 109 RFU for FLASH and 870 ± 51 RFU for CONV at 10 Gy (p < 0.001) (**Fig. 2b**). Notably, LPA-FLASH-RT almost completely protected normal BEAS-2B cells from this damage, keeping the intensity at 163 ± 15 RFU. In contrast, CONV-RT caused normal cells to suffer high lipid damage, reaching 714 ±74 RFU at 10 Gy (p < 0.001). These results show that LPA-FLASH-RT specifically shields healthy cell membranes from oxidative insult.

Since LPO is a major part of iron-dependent catalysis, we looked at changes in the labile iron pool (Fe^2+^). Qualitative images using the FerroFarRed probe showed bright red signals in A549 tumor cells at 5 Gy for both radiation types, which means high iron accumulation (**Fig. 2c**). In normal BEAS-2B cells, the LPA-FLASH-RT group showed very little red signal, but the CONV-RT group showed a clear increase in iron release (**Fig. 2c**). Quantitative measurements confirmed these patterns (**Fig. 2d**). In A549 cancer cells at 10 Gy, iron levels reached 15.6 ± 1.2 MFI after LPA-FLASH-RT and 14.6 ± 2.3 MFI after CONV-RT (p < 0.001). For normal BEAS-2B cells, LPA-FLASH-RT effectively stopped iron overload, with levels staying low at 5.6 ± 1.1 MFI compared with 11.6 ± 1.8 MFI seen under CONV-RT at 10 Gy (p < 0.001). Together, these findings show that the protection of normal tissue by laser-driven FLASH protons comes from blocking a chain of oxidative stress, iron release, and lipid damage.

To understand the relationship between iron accumulation and membrane damage, we analyzed the correlation between intracellular labile iron (Fe^2+^) and lipid peroxidation (LPO) levels. We observed a strong positive correlation between these two markers across both cell lines (**Fig. 2e**). Normal BEAS-2B cells treated with 5 Gy LPA-FLASH-RT remained in the low-iron and low-LPO region, while those treated with 5 Gy CONV-RT shifted significantly toward the high-damage region. These results indicating that the FLASH effect in healthy tissue is rooted in the interruption of the oxidative-iron-lipid damage cascade.

To decipher the molecular “gatekeeper” responsible for this selective protection within the 1–4 hour window, we focused on the stress-responsive transcription factor ATF3, which we previously identified as a key signature of the FLASH effect [**13**]. We proposed a molecular model where the nanosecond pulse probes a transcriptional blind spot, effectively bypassing the ATF3-mediated suppression of the SLC7A11 system (**Fig. 2f**). To test this model, we qualitatively assessed the expression of ATF3 at 4 hours post-irradiation using immunofluorescence imaging. In A549 tumor cells, both LPA-FLASH-RT and CONV-RT at 5 Gy induced strong red signals in the nuclei, indicating high ATF3 activation (**Fig. 2g**). In normal BEAS-2B cells, however, we observed a clear difference: CONV-RT triggered high ATF3 expression, while LPA-FLASH-RT resulted in almost no detectable red signal.

Quantitative analysis of ATF3 expression levels confirmed these observations. In A549 tumor cells, ATF3 increased with the radiation dose, reaching a mean fluorescence intensity (MFI) of 126.1 ± 8.2 for FLASH and 124.0 ± 10.5 for CONV at 10 Gy (p < 0.001 relative to 0 Gy; **Fig. 2h**). In contrast, normal BEAS-2B cells treated with LPA-FLASH-RT remained at near-baseline levels, reaching only 13.8 ± 1.5 MFI at the highest dose. This protection was absent under conventional radiation, where CONV-RT induced a significant rise in ATF3 for normal cells, reaching 98.5 ± 15.1 MFI at 10 Gy (p < 0.001; **Fig. 2h**). Together, these results demonstrate that the temporal singularity of LPA-FLASH-RT enables normal cells to remain beneath the kinetic threshold for ATF3 activation, thereby bypassing the pro-ferroptotic axis and safeguarding the cellular antioxidant infrastructure.

### Selective preservation of mitochondrial cristae and bioenergetic efficiencty under LPA-FLASH irradiation

Since ATF3 acts as a master regulator of mitochondrial iron homeostasis and antioxidant defense, we investigated whether its selective suppression in FLASH-irradiated normal cells translates into preserved mitochondrial bioenergetics. Beyond mere survival, we sought to determine if the avoidance of the ATF3-ferroptotic axis enables a state of mitochondrial hormesis, where healthy cells maintain or even enhance their metabolic output under radiation stress. We first quantified intracellular ATP levels at 4 h post-irradiation to assess the metabolic state of the cells (**Fig. 3a**). In A549 tumor cells, both irradiation modalities at 5 Gy induced a significant bioenergetic decline. Specifically, ATP content decreased from 6.6 ± 0.2 RFU to 5.0 ± 0.1 RFU following LPA-FLASH-RT and to 5.6 ± 0.3 RFU following CONV-RT (p<0.01 for both). In striking contrast, normal BEAS-2B cells exhibited a divergent metabolic response. While conventional CONV-RT resulted in a substantial reduction of ATP levels to 6.2 ± 0.4 RFU (p<0.01), LPA-FLASH-RT treatment prompted an unexpected metabolic surge, elevating ATP levels to 9.4 ± 0.2 RFU (p<0.01). This suggests that ultra-high-dose-rate protons may trigger a compensatory bioenergetic response or maintain higher mitochondrial efficiency in healthy tissue.

**Figure 3.**
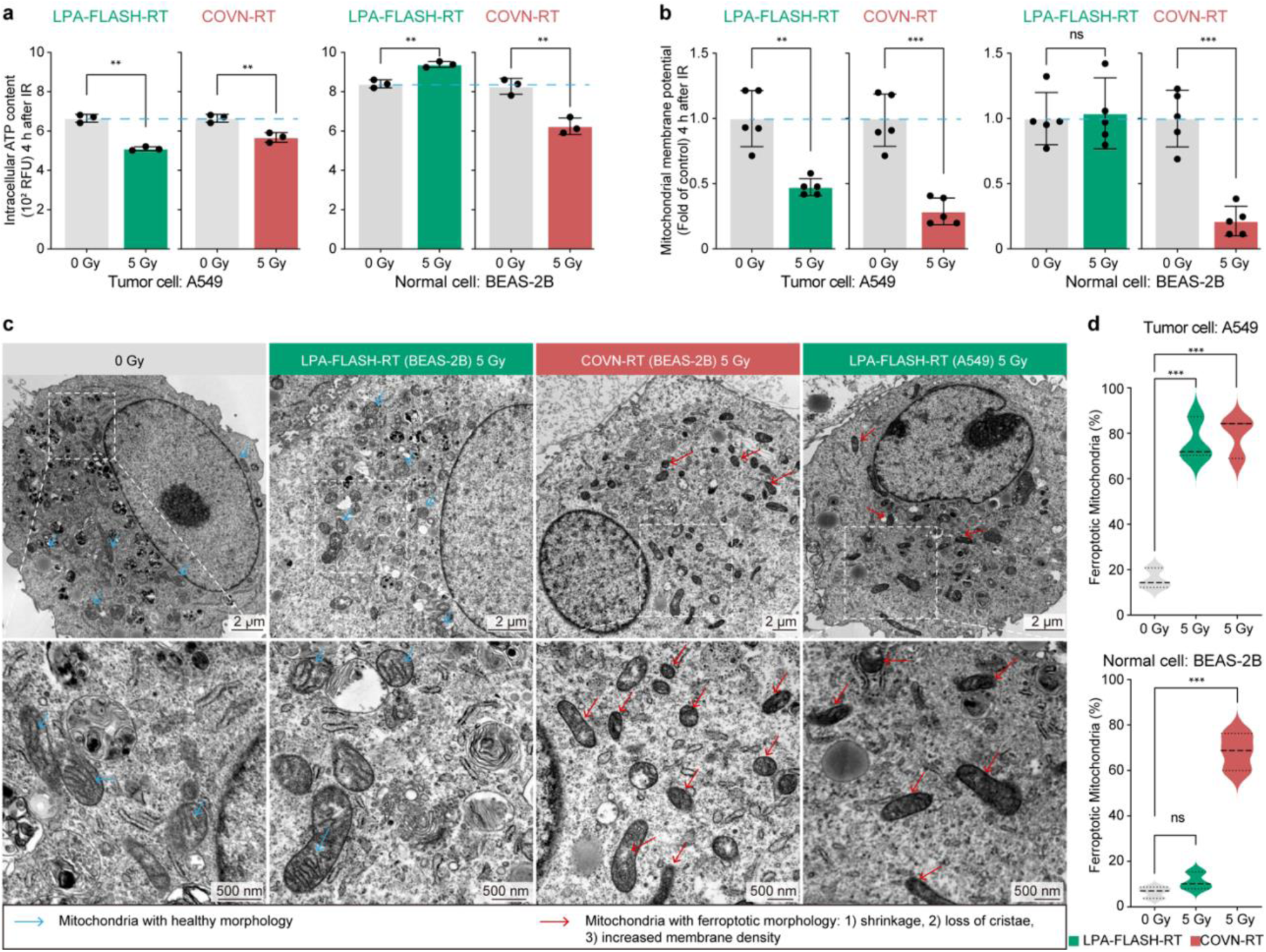
FLASH irradiation preserves mitochondrial bioenergetics and ultrastructural integrity in normal tissues. **(a, b)** Assessment of mitochondrial function in BEAS-2B and A549 cells at **4 h** post-irradiation. Intracellular ATP levels **(a)** and the mitochondrial membrane potential **(b)**, indicated by the ratio of red (aggregates) to green (monomers) JC-1 fluorescence, were normalized to non-irradiated controls. **(c)** Representative transmission electron microscopy (TEM) images showing mitochondrial morphology. The upper panels display low-magnification overviews (Scale bar, 20 μm), and the lower panels show high-magnification insets of the selected regions (Scale bar, 500 nm). Blue arrows indicate healthy mitochondria with intact cristae, while red arrows indicate mitochondria with characteristic ferroptotic features (shrinkage, membrane densification, and cristae loss). **(d)** Quantification of the percentage of mitochondria exhibiting ferroptotic morphology, presented as violin plots. The upper panel shows results for A549 tumor cells, and the lower panel for BEAS-2B normal cells. Green distributions represent LPA-FLASH-RT (5 Gy), and red distributions represent CONV-RT (5 Gy). Data are presented as mean ± s.d. for **(a, b)** (n=3 independent experiments) and as distribution density for (**d**) (n=3 random fields). Statistical significance was analyzed using one-way ANOVA; ns, no significant difference (p > 0.05); *, significant difference (p < 0.05); **, very significant difference (p < 0.01); ***, highly significant difference (p < 0.001).

We further examined the mitochondrial membrane potential (MMP) as a critical indicator of organelle health (**Fig. 3b**). Malignant A549 cells showed a sharp dissipation of MMP under both modalities, with potential dropping to 0.46 ± 0.05 (FLASH-RT) and 0.28 ± 0.1 (CONV-RT) of the control baselines (p<0.01 and p < 0.001, respectively). For normal BEAS-2B cells, CONV-RT led to a severe mitochondrial collapse (0.2 ± 0.1 of control, p < 0.001), whereas LPA-FLASH-RT effectively maintained the potential at 1.03 ± 0.3, showing no statistically significant difference from the unirradiated control (p > 0.05).

To determine if the observed functional bioenergetic failures were accompanied by structural alterations, we examined mitochondrial ultrastructure using transmission electron microscopy (TEM) at 4 h post-irradiation (**Fig. 3c**). In unirradiated control cells, mitochondria typically exhibited a healthy morphology, characterized by distinct, well-organized cristae and moderate membrane density (blue arrows). However, following irradiation, we observed a shift toward a “ferroptotic morphology” in specific groups, defined by mitochondrial shrinkage, loss or rupture of cristae, and significantly increased outer membrane density (red arrows).

Qualitative TEM analysis revealed that in A549 tumor cells, both LPA-FLASH-RT and CONV-RT induced severe structural degradation at 5 Gy, with the majority of mitochondria appearing shrunken and devoid of internal cristae structures (**Fig. 3c**). In striking contrast, the mitochondrial ultrastructure of normal BEAS-2B cells remained largely preserved under LPA-FLASH-RT, maintaining clear cristae similar to the control group. Conversely, BEAS-2B cells subjected to CONV-RT underwent widespread mitochondrial collapse, displaying the same characteristic ferroptotic shrinkage and membrane thickening observed in the tumor cell lines (**Fig. 3c**).

Quantitative assessment of the percentage of ferroptotic mitochondria confirmed these morphological observations (**Fig. 3d**). In malignant A549 cells, the proportion of ferroptotic mitochondria increased significantly from a baseline of approximately 14% to 72.1 ± 8.5% under LPA-FLASH-RT and 84.5 ± 7.2% under CONV-RT (p < 0.001 for both). For normal BEAS-2B cells, the impact was highly dependent on the dose rate. While conventional radiation (CONV-RT) triggered a massive structural shift, with 68.4±9.1% of mitochondria showing ferroptotic features (p < 0.001), LPA-FLASH-RT maintained the percentage of damaged mitochondria at a near-baseline level of 10.2 ± 3.4%, which was not statistically significant compared to the control group (p > 0.05).

Combined with the previously identified functional preservation of ATP and membrane potential, these ultrastructural findings provide definitive evidence that laser-driven FLASH protons selectively safeguard mitochondrial architecture, preventing the functional collapse of the cell’s metabolic engine.

### Volumetric 3D imaging confirms the structural resilience of the global mitochondrial reticulum in FLASH-irradiated normal cells

The preservation of mitochondrial cristae observed via TEM provided a snapshot of structural resilience; however, to confirm that this protection extends to the entire cellular metabolic infrastructure, we performed whole-cell 3D light-sheet microscopy (**Fig. 4a**). This approach allowed for the precise quantification of mitochondrial spatial organization and morphological complexity across the entire cellular volume, providing a volumetric validation of the laser-driven FLASH protons induced sparing effect.

**Fig. 4.**
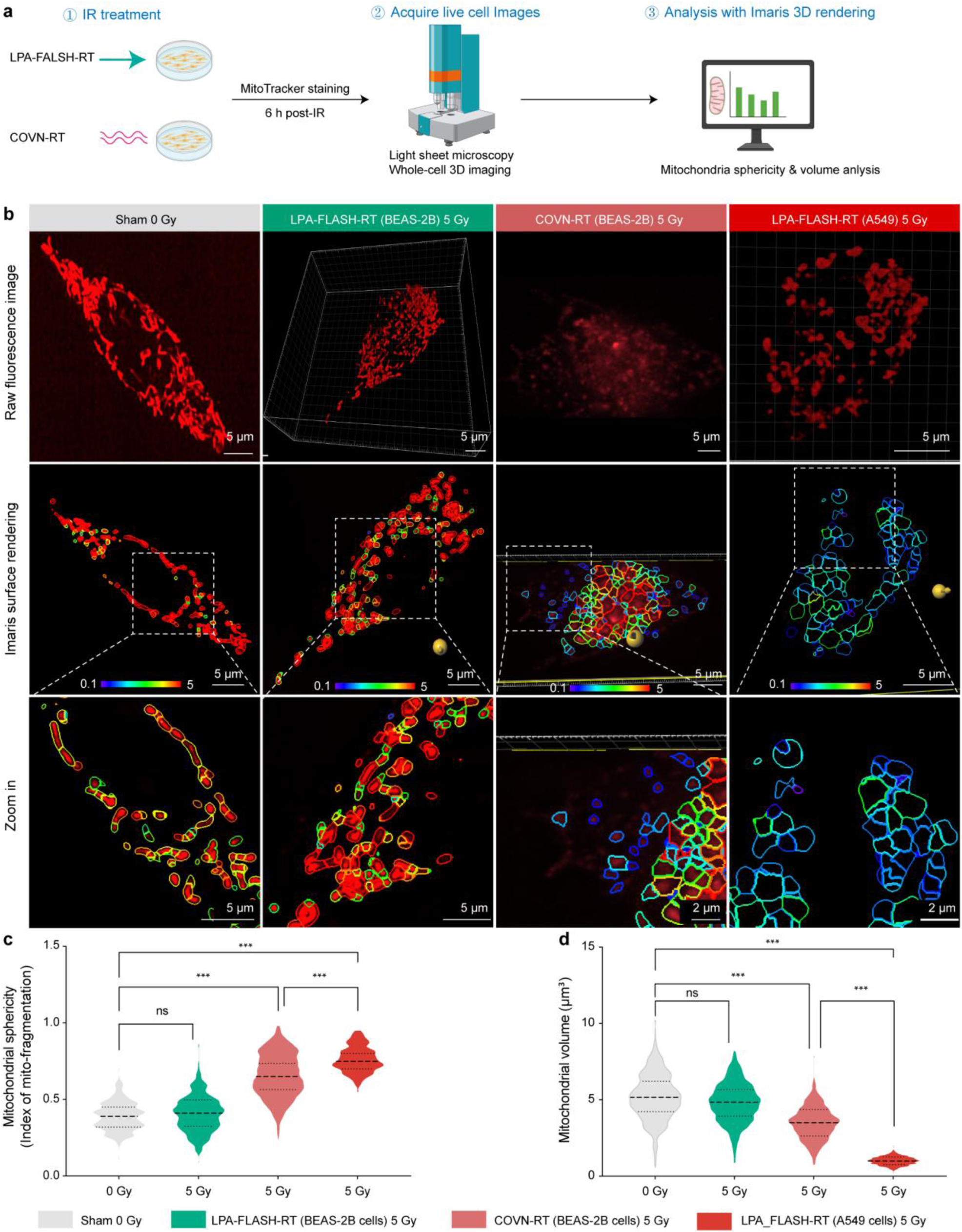
FLASH irradiation preserves mitochondrial network topology and volumetric homeostasis. a, Schematic representation of the experimental and computational pipeline. **a,** Workflow for cell morphometric quantification by light-sheet fluorescence microscopy. Normal (BEAS-2B) and malignant (A549) lung cells were subjected to either LPA-FLASH-RT or CONV-RT (5 Gy). Sub-cellular mitochondrial architecture was captured via high-resolution 3D light-sheet fluorescence microscopy (LSFM), followed by 3D surface rendering and morphometric quantification (sphericity and volume) using Imaris software. **b,** Representative images of mitochondrial morphology across treatment groups. Top, raw fluorescence signals; middle, corresponding 3D surface-rendered models; bottom, magnified views of the regions indicated by white dashed boxes. Scale bars, 5 μm; zoom-in scale bars, 5 μm and 2 μm. **c, d,** Quantitative analysis of mitochondrial sphericity (c) and mean mitochondrial volume (d). Sphericity values approaching 1.0 indicate a fragmented, spherical state. Volume data reflect radiation-induced mitochondrial swelling (BEAS-2B) or degradation (A549). Data are presented as mean ± s.d. from n=600 independent biological replicates. Statistical significance was determined by one-way ANOVA with Tukey’s post-hoc test for multiple comparisons. ns, no significant difference (p > 0.05); *, significant difference (p < 0.05); **, very significant difference (p < 0.01); ***, highly significant difference (p < 0.001).

Qualitative 3D reconstruction revealed that mitochondria in unirradiated control cells and normal BEAS-2B cells subjected to LPA-FLASH-RT maintained a highly interconnected, elongated, and tubular reticulum (**Fig. 4b**). In striking contrast, BEAS-2B cells irradiated with conventional protons (CONV-RT) exhibited a fragmented phenotype, with the network collapsing into discrete, punctate structures. This morphological degradation was most severe in malignant A549 cells under LPA-FLASH-RT, which demonstrated extreme mitochondrial fragmentation and clustering.

To quantify these architectural shifts, we calculated the mitochondrial sphericity—serving as an index of fragmentation where higher values represent more rounded, isolated mitochondria—and the total mitochondrial volume per cell (**Fig. 4c, d**). In normal BEAS-2B cells, LPA-FLASH-RT successfully preserved the mitochondrial architecture; the sphericity index (0.45 ± 0.08) and mitochondrial volume (4.9 ± 1.1 μm^3^) showed no significant deviation from the sham-irradiated control (p > 0.05; **Fig. 4c, d**). Conversely, CONV-RT triggered significant network disruption in normal cells, as evidenced by a marked increase in sphericity to 0.65 ± 0.12 (p < 0.001) and a reduction in volume to 3.6 ± 0.9 μm^3^ (p < 0.001). The most dramatic failure was observed in tumor A549 cells under LPA-FLASH-RT, where sphericity escalated to 0.75 ± 0.10 and volume plummeted to 1.0 ± 0.4 μm^3^ (p < 0.001 for both). These whole-cell imaging data corroborate our ultrastructural findings and confirm that laser-driven FLASH protons selectively safeguard 3D mitochondrial network integrity in normal tissue while inducing catastrophic morphological and functional failure in malignant cells.

## Discussion

The therapeutic promise of the FLASH effect has been constrained by the limitation of conventional accelerators and a fragmented understanding of its underlying radiobiological mechanisms. In this study, by utilizing the 200-TW beamline at the SIOM HLDP platform, we demonstrated a special “temporal singularity” effect of nanosecond laser-driven proton FLASH, where proton doses were delivered via discrete 12.9 ns pulses at an extreme instantaneous rate of 1.94×10^7^ Gy/s. This effect, as elucidated in the 2026 Nature Reviews Cancer (NRC) analysis by Vozenin et al., suggests the paradigm which shifts from a narrow focus on radiolytic oxygen depletion (ROD) toward an integrated framework of physicochemical radical kinetics [**16**]. In this effect, the nanosecond proton delivery induces a “biological decoupling” phenomenon, where the radical-radical recombination is much faster than the activation kinetics of cellular stress-sensing pathways. This mechanism physically reduces the effective yield of oxidative products before they can interact with the biological milieu [**17**].

While the previous studies of the FLASH effect focused on the attribution of rapid oxygen depletion [**8**], recent evidence suggests that ferroptosis—an iron-dependent form of regulated cell death—may be a pivotal mediator in the differential response between tumor and normal tissues [**9,10**]. Specifically, ionizing radiation is known to trigger ferroptosis by elevating ROS and upregulating ACSL4 [**18–20**]. Therefore, in this study, we hypothesized the execution of FLASH-induced ferroptosis for the primary “biological decoupling”effect, and indeed, we have identified ATF3 as a central regulatory hub in this process. In the FLASH treatment, transcriptional repression of SLC7A11 works as a temporal sensor that critically distinguishes the FLASH-RT response from CONV-RT, thereby sensitizing tumor cells to ultra-high-dose-rate irradiation. Under CONV-RT, the sustained accumulation of reactive oxygen species (ROS) triggers the transcriptional activation of ATF3, which subsequently represses the SLC7A11/xCT antioxidant system, mobilizing the mitochondrial labile iron pool and catalyzing lethal lipid peroxidation. In contrast, the FLASH pulse appears to deliver the dose within a temporal “blind spot” for ATF3 activation in normal cells, effectively maintaining the cell’s antioxidant infrastructure and preventing the transition to ferroptotic cell death.

A vital question addressed by this work is why this protection “FLASH effect” does not extend to malignant A549 cells. We posited that this selectivity is rooted in the pre-existing metabolic divergence between healthy and cancerous tissues [**21**]. Tumor cells exist in a state of primed oxidative stress and iron metabolic dysregulation, which significantly lowers their threshold for ferroptotic execution. Consequently, the unique temporal structure of a FLASH pulse—while sparing for homeostatic tissue—remains insufficient to prevent the catastrophic failure of their already-strained antioxidant defenses in malignant cells. In contrast, normal BEAS-2B cells leverage the transcriptional blind spot to mount a robust adaptive resilience. Mechanistically, by bypassing the ATF3-SLC7A11 transcriptional blockade, normal cells prevent the ferroptotic chain reaction that typically leads to mitochondrial collapse. Indeed, in this work, we for the first directly observed the remarkable preservation of mitochondria cristae integrity—quantified via TEM and 3D light-sheet imaging, confirming that laser-driven FLASH protons protect the very bioenergetic cores of the cell. In this protected state, the mitochondrial reticulum remains highly interconnected and elongated, providing the structural scaffold necessary for efficient cellular respiration. Particularly, we also observed the associated ATP surge during the temperal process, and we interpret this ATP surge not merely as a lack of damage, but as an active compensatory response. This functional resilience is mirrored by the dramatic nine-fold reduction in the α value, suggesting that laser-driven FLASH protons enable healthy tissue to maintain, or even temporarily enhance, its metabolic output under radiation stress.

Furthermore, our results confirm that ferroptosis, rather than traditional apoptosis or necrosis, serves as the primary cell death pathway modulated by the FLASH effect in this nanosecond laser-proton regime. This claim offers an intriguing contrast to the recent work by Yan et al.[**22**], who demonstrated that electron-driven FLASH-RT primarily regulates mitochondrial Cytochrome c leakage and subsequent caspase-mediated apoptosis. We proposed that this shift in execution pathways is fundamentally governed by the energy deposition density (Δ*E*). While electron beams are characterized by diffuse ionization and lower Linear Energy Transfer (LET)—with Geant4 Monte Carlo simulations showing ∼1 keV in a 5 μm cell layer—our laser-accelerated protons deliver extreme instantaneous energy within localized volumes. According to our simulations, 7 MeV LPA-FLASH-RT protons traversing a 3.6 μm Mylar foil exhibit a peak energy deposition of 55 keV within a 5 μm cellular surrogate, corresponding to an effective LET of ∼11 keV/μm. In comparison, 221 MeV clinical-energy protons yield a peak deposition of only ∼13 keV (∼2.6 keV/μm), indicating an approximately 4-fold increase in local energy deposition, which allows the effective LET of FLASH protons to approach to that of therapeutic heavy ions (for example, the LET in the plateau region of therapeutic carbon-ion beams is typically on the order of tens of keV/μm [**23]**). Moreover, both experimental measurements and simulations indicate that energy degradation in the solid-water phantom substantially reduces the number of protons reaching the cellular layer in the CONV-RT configuration, while simultaneously broadening the incident energy spectrum and the distribution of deposited energies. In contrast, FLASH-RT produces a relatively narrow incident energy spectrum, leading to more uniform energy deposition within the cellular layer (**Supplementary Content 1 and Supplementary Fig. 1**). The magnitude of Δ*E* serves as a physical switch for cell fate. The extreme Δ*E* of laser-protons induces immediate and catastrophic iron-dependent lipid peroxidation, while the lower Δ*E* of electrons triggers organized, protein-mediated signaling—the classic apoptotic cascade. This explains the difference against electron-driven FLASH-RT, and also suggests that the biological FLASH effect is not merely a function of the average high dose rate, but is intrinsically tied to the discrete spatial-temporal distribution of energy at the sub-cellular level.

In summary, this study delineates the mechanistic advancements of the FLASH effect across some more critical dimensions, marking a conceptual transition from passive physical shielding to active metabolic orchestration (**Fig.5**). First, at the temporal limit, we establish that LPA-FLASH-RT—operating at a 12.9 ns pulse scale—induces a temporal singularity where ultra-fast radiolytic chemistry is effectively decoupled from the kinetic constraints of biological stress sensing. Second, we especially identify ATF3 as a pivotal molecular temporal sensor, revealing that the nanosecond-scale delivery probes a “transcriptional blind spot” that allows normal cells to remain beneath the kinetic threshold required for stress-responsive activation. Third, our work ambiguously confirms that the ferroptotic cascade—driven by mitochondrial labile iron mobilization rather than conventional apoptotic pathways—serves as the primary executioner of the FLASH therapeutic window. Fourth, this work provides the direct evidence of the preservation of mitochondrial cristae integrity and the subsequent functional ATP surge, so that we define metabolic resilience as the fundamental determinant of cellular survival under extreme physical stress. Beyond these mechanistic insights, laser-driven proton sources offer transformative advantages over traditional radiofrequency accelerators, including their inherent compactness, potential for lower facility footprints, and the unique capability to deliver extreme instantaneous dose rates through nanosecond-scale pulses.

**Fig. 5.**
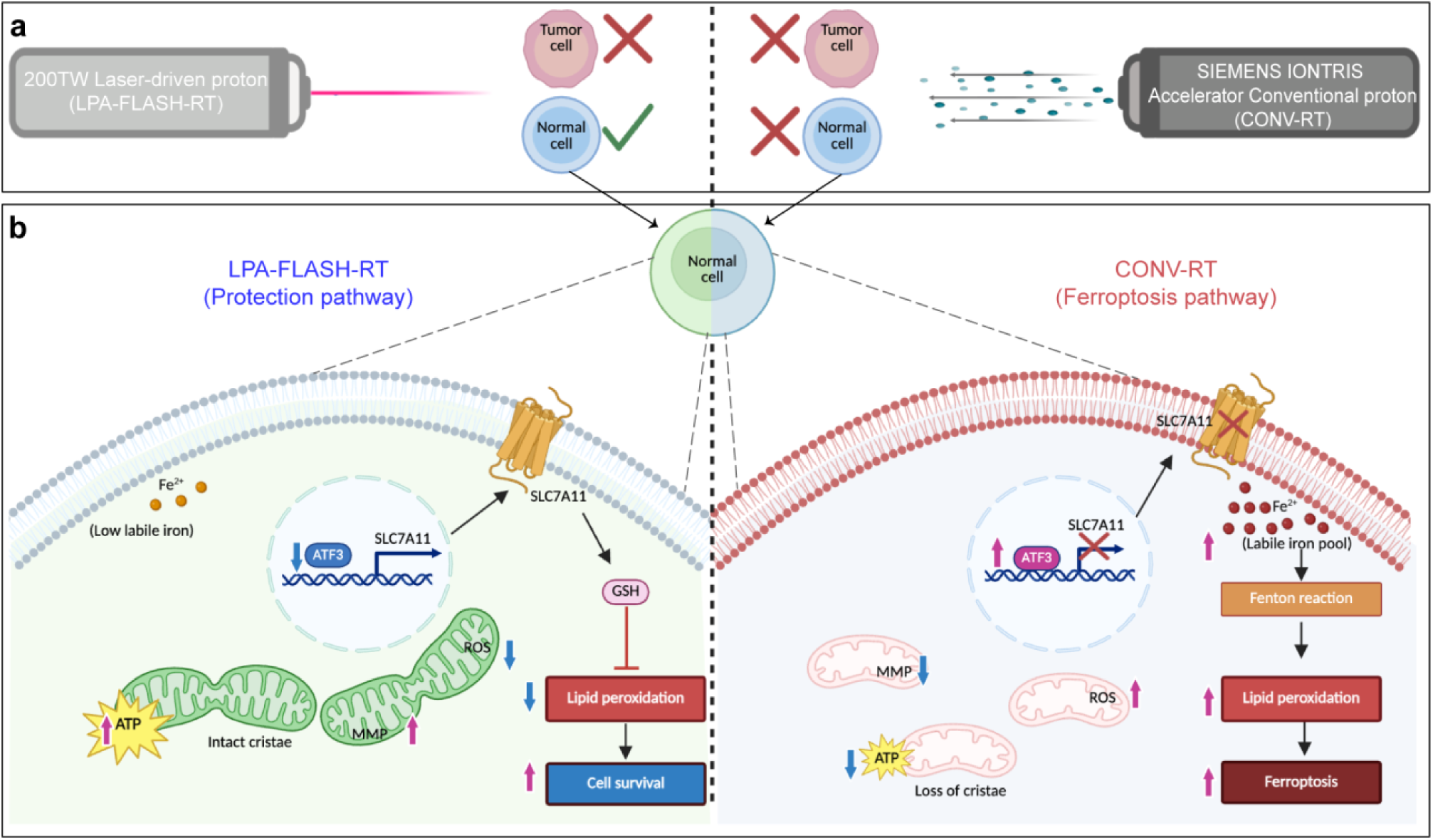
Schematic mechanism of laser-driven ultra-high dose-rate proton radiotherapy selectively sparing normal lung cells via the ATF3-ferroptosis axis. **a,** Macro-level irradiation patterns and cellular survival. FLASH proton beams driven by a petawatt-class laser, characterized by ultra-short pulses and extreme instantaneous dose rates (>10^7^ Gy/s), are shown on the left, while conventional proton (CONV-RT) irradiation with continuous low-intensity beams is illustrated on the right at equivalent doses, LPA-FLASH-RT effectively kills A549 tumor cells while significantly enhancing the clonogenic survival of BEAS-2B normal epithelial cells, demonstrating a pronounced FLASH effect. **b,** the molecular interplay governing the differential fate of normal cells. Left panel (LPA-FLASH-RT protection pathway): The ultra-high dose rate maintains a low labile iron pool (Fe^2+^) and suppresses ATF3 expression in the nucleus. This allows for continued transcription and function of the SLC7A11 transporter, promoting glutathione (GSH) synthesis. GSH inhibits reactive oxygen species (ROS) accumulation and lipid peroxidation, thereby preserving mitochondrial ultrastructure (intact cristae), maintaining high mitochondrial membrane potential (MMP) and ATP production, ultimately favoring cell survival. Right panel (CONV-RT ferroptosis pathway): Conventional irradiation leads to an expanded labile iron pool and upregulation of ATF3, which transcriptionally represses SLC7A11. The consequent failure of the antioxidant system drives iron-dependent Fenton reactions, leading to severe lipid peroxidation, mitochondrial dysfunction (loss of cristae, reduced MMP and ATP), and the execution of ferroptosis. Up arrows (↑) indicate upregulation or increase; down arrows (↓) indicate downregulation or decrease; crossed gene indicates transcriptional repression; crossed transporter indicates functional loss.

Looking forward, these findings collectively advocate for a paradigm shift in radiation biology: from a traditional DNA-centric model to a metabolic-driven framework of cell fate determination. The elucidation of the role of ATF3-mitochondrial iron axis in FLASH-RT suggests a future of biomarker-guided radiotherapy, where the therapeutic index can be rationally optimized by modulating cellular ferroptotic thresholds, and thus may provide the scientific foundation for the rational design of next-generation, high-precision radiotherapy platforms.

## Method

### 1. Cell lines and culture conditions

Human lung adenocarcinoma cells (A549) and human normal bronchial epithelial cells (BEAS-2B) were obtained from the American Type Culture Collection (ATCC). Cells were maintained in DMEM (Gibco) supplemented with 10% fetal bovine serum (FBS) and 1% penicillin-streptomycin in a humidified incubator at 37°C with 5% CO_2_. For all irradiation experiments, cells were seeded onto custom-designed culture dishes sealed with a 2 μm Mylar foil base. This thin interface was critical to minimize proton energy loss and ensure accurate dose delivery to the cell monolayer.

### 2. Proton irradiation and dosimetry

#### Laser-driven FLASH-RT

Ultra-high-dose-rate irradiation was performed using a 200 TW Laser System. Protons were accelerated to 7 MeV and delivered in pulses (repetition rate: 0.1 Hz; pulse width: 12.9 ns). The system achieved an extreme instantaneous dose rate of 1.94×10^7^ Gy/s. Cells were irradiated within a vacuum-compatible chamber, with the beam passing through a vacuum window before reaching the Mylar foil interface.

#### Conventional-RT (CONV-RT)

Standard dose-rate irradiation was conducted using a SIEMENS IONTRIS Accelerator System. A proton beam with an initial energy of 221 MeV was degraded using a solid water phantom so that the protons reached the Bragg peak region at the position of the cell monolayer, corresponding to proton energies in the few-MeV range after passing through the beamline materials. The dose rate was maintained at a steady-state 0.08 Gy/s.

#### Dosimetry

All doses (2, 5, and 10 Gy) were calibrated and verified using EBT3 radiochromic films (Gafchromic). Absolute dose accuracy was maintained within ± 5% across both modalities. Detailed descriptions of the irradiation geometry and the corresponding Geant4 Monte Carlo simulations of proton energy deposition in the cell layer are provided in Supplementary information Content 1 and Supplementary Fig. 1 and Supplementary Fig. 2.

### 3. Clonogenic survival assay and LQ modeling

Following irradiation, cells were immediately trypsinized and re-seeded into 6-well plates at densities optimized for each dose. After 10–14 days of incubation, colonies were fixed with methanol and stained with 0.5% crystal violet. Colonies consisting of >50 cells were counted manually. The survival fraction (SF) was calculated and fitted using the linear-quadratic (LQ) model: S=e−^(αD+βD2)^, where D is the dose and α, β are the radiosensitivity coefficients.

### 4. Immunofluorescence staining

Cells were fixed with 4% paraformaldehyde (PFA) for 15 min and permeabilized with 0.5% Triton X-100. After blocking with 5% BSA, samples were incubated overnight at 4°C with primary antibodies against γ-H2AX (1:500, Proteintech) or ATF3 (1:200, Proteintech). Cells were then labeled with fluorophore-conjugated secondary antibodies and counterstained with DAPI. Images were acquired using a Leica SP8 confocal microscope. For γ-H2AX, DNA double-strand breaks (DSBs) were quantified as the number of foci per nucleus. For ATF3, the mean fluorescence intensity (MFI) within the nucleus was calculated using ImageJ.

### 5. Assessment of oxidative stress and ferroptosis markers

ROS and Lipid Peroxidation: Intracellular reactive oxygen species (ROS) were measured 1 h post-irradiation using the H_2_DCFDA probe (10 μM). ROS levels were quantified via flow cytometry (BD LSRFortessa). Lipid peroxidation (LPO) was assessed 4 h post-irradiation using the Liperfluo probe (1 μM). LPO-specific fluorescence was captured via confocal microscopy and analyzed as relative fluorescence units (RFU).

Labile Iron Pool (Fe^2+^): Intracellular ferrous iron levels were evaluated 4 h post-irradiation using the FerroFarRedfluorescent probe (1 μM, Goryo Chemical). After 30 min of incubation at 37°C, cells were imaged using confocal microscopy (Ex/Em: 635/660 nm). Intensity was quantified as MFI to assess the correlation between iron overload and lipid damage.

### 6. Mitochondrial functional and structural analysis

ATP and MMP: Intracellular ATP was quantified 4 h post-irradiation using a luminescence-based assay kit (Beyotime) and normalized to total protein concentration. Mitochondrial membrane potential (MMP) was determined using the JC-1probe. The ratio of red (aggregates) to green (monomers) fluorescence was calculated to evaluate mitochondrial polarization.

Transmission Electron Microscopy (TEM): Cells were fixed in 2.5% glutaraldehyde and post-fixed in 1% osmium tetroxide. Samples were dehydrated and embedded in epoxy resin. Ultrathin sections (70–90 nm) were stained with uranyl acetate and lead citrate and imaged using a Hitachi H-7650 TEM at 80 kV. Mitochondria were classified as ferroptotic based on characteristic shrinkage, loss of cristae, and increased membrane density.

### 7. 3D Mitochondrial network dynamics

For whole-cell morphological analysis, cells were stained with MitoTracker Deep Red FM. Z-stack images were acquired using a light-sheet fluorescence microscope (Z-step: 100 nm). 3D reconstruction and surface rendering were performed using Imaris 10.1. Mitochondrial sphericity (an index of fragmentation) and total volume were quantified across the entire cell volume. Network skeletonization was performed using the MiNA plugin in ImageJ to calculate branching and complexity.

### 8. Statistical analysis

All experiments were performed in at least three independent biological replicates. Data are presented as mean ± s.d. Statistical significance was evaluated using one-way ANOVA followed by Tukey’s *post-hoc* test for multiple comparisons or Student’s *t*-test for pairwise comparisons. P<0.05 was considered statistically significant (*, p < 0.05; **, p < 0.01; ***, p < 0.001). All analyses were conducted using GraphPad Prism 10.2.

## Acknowledgements

This work was supported by the National Key R&D Program of China (Grant No. 2025YFF0515103), the National Natural Science Foundation of China (Grant No. 42225405), the CAS Project for Young Scientists in Basic Research (Grant No. YSBR060), the Innovation Fund of the Anti-Radiation Application Technology Innovation Center of China Institute of Atomic Energy (Grant No. KFZC2023021002), and the Interdisciplinary Cultivation Program at Shandong University, Weihai.

## Supplementary information

Supplementary information is available on-line for this paper.

---

**Supplementary Fig 1.**
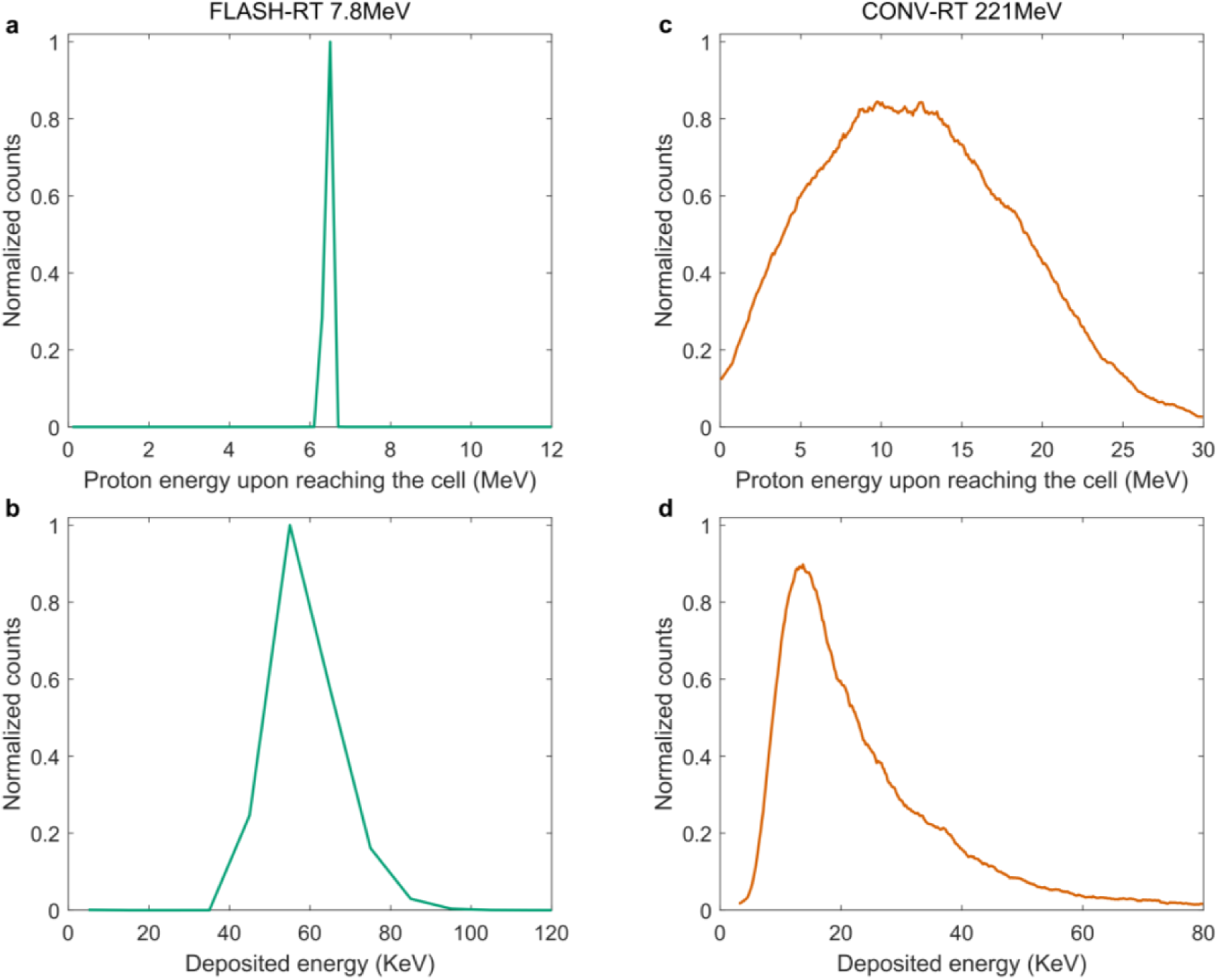
Geant4-simulated energy deposition spectra of protons under representative irradiation conditions. **a**, Incident proton energy spectrum before the cellular layer, with a peak at approximately 6.5 MeV. **b,** Corresponding energy deposition spectrum in the 5 μm aqueous layer for the incident spectrum in (a), exhibiting a peak at approximately 55 keV. **c,** Incident proton energy spectrum for 221 MeV protons after traversing the composite beamline, showing a relatively broad distribution with a main peak at 8-14 MeV. **d,** Corresponding energy deposition spectrum in the 5 μm aqueous layer for the incident spectrum in (c), peaking at approximately 13 keV. The green curves represent LPA-FLASH-RT data, and the red curves represent CONV-RT data. For both configurations, each spectrum was generated using a primary particle count of 1× 10^7^ protons.

**Supplementary Fig. 2.**
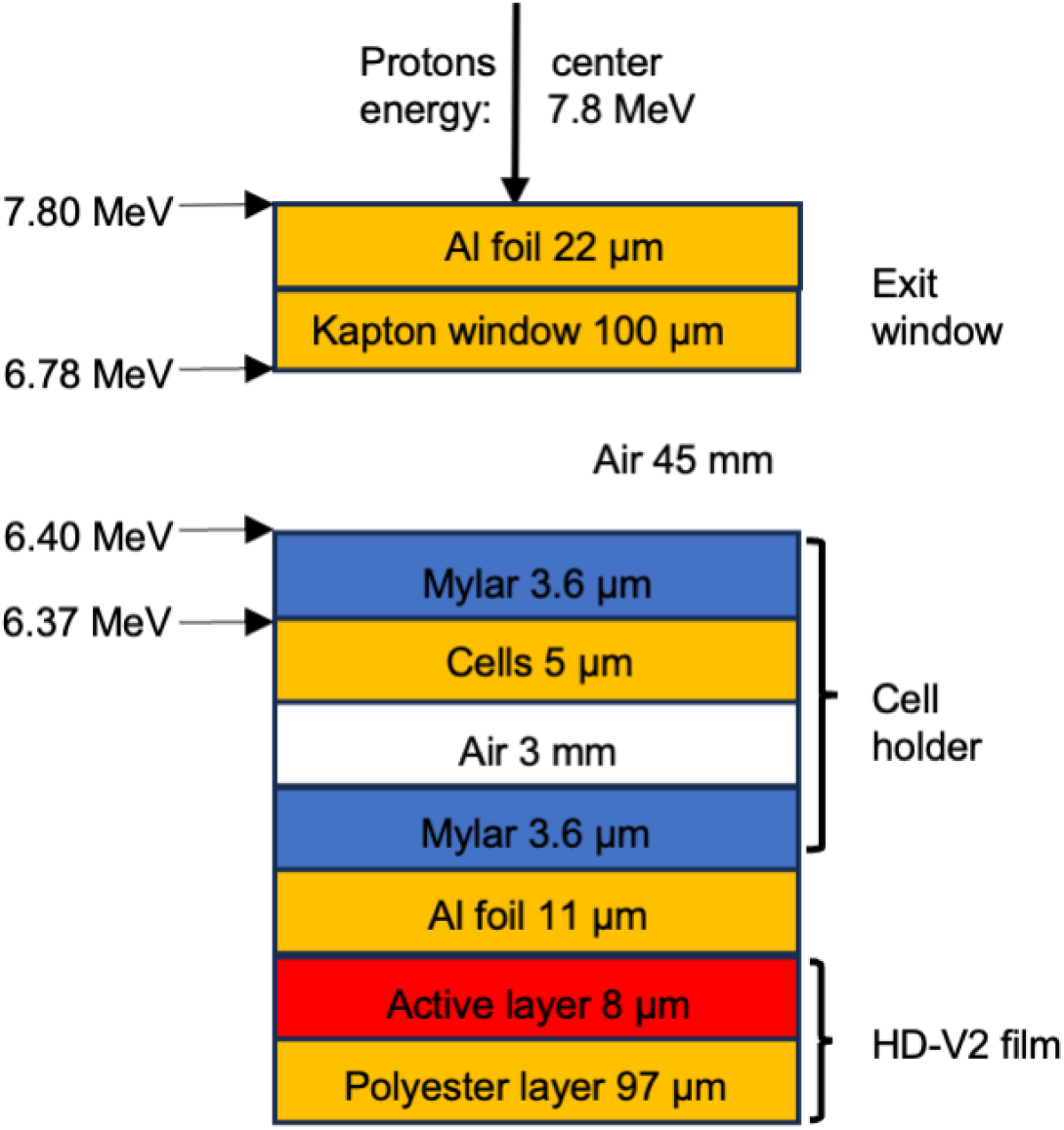
Schematic and calibration of the proton dose delivery system.

