## Supplementary material for "Nanosecond laser-driven proton FLASH spares normal tissue cells by sustaining mitochondrial homeostasis and attenuating ferroptosis": SI

**CONTENTS**

**1. GEANT4 Monte Carlo simulation of proton energy deposition spectra in the cell layer under LPA-FLASH-RT and CONV-RT**

**2. Dosimetry**

1. **GEANT4 Monte Carlo simulation of proton energy deposition spectra in the cell layer under LPA-FLASH-RT and CONV-RT**

Geant4 Monte Carlo simulations were employed to model the energy deposition spectra of protons under configurations representative of LPA-FLASH-RT and conventional radiotherapy (CONV-RT). In the configuration for (a) and (b), 7.8 MeV protons traversed a 22 μm aluminum foil, a 100 μm Kapton window, and 45 mm of air before reaching a 3.6 μm Mylar foil (C_10_H_8_O_4_, ρ = 1.3 g/cm^3^), followed by a 5 μm aqueous layer serving as a cellular surrogate. The incident proton energy spectrum before the cellular layer exhibits a peak at approximately 6.5 MeV (Fig. S1a), while the corresponding energy deposition spectrum in the aqueous layer peaks at about 55 keV (Fig. S1b). In the configuration for (c) and (d), 221 MeV protons traversed a composite beamline—comprising 80 cm of air, 30 cm of solid water (98% C_8_H_8_ + 2% TiO_2_, ρ = 1.045 g/cm^3^), 10 cm of air, and 0.3 mm of polyvinyl chloride (PVC, C_2_H_3_Cl, ρ = 1.4 g/cm^3^)—prior to reaching the 5 μm water target. The thickness of the solid water phantom was chosen so that the Bragg peak occurred close to the cellular layer after accounting for energy losses in the upstream air gaps and beamline materials. Under this configuration, the incident proton energy spectrum before the cellular layer exhibits a relatively broad distribution with a main peak in the range of 8-14 MeV (Fig. S1c), while the corresponding energy deposition spectrum in the aqueous layer peaks at about 13 keV (Fig. S1d). Each simulation utilized a primary particle count of 1× 10^7^ protons. The results indicate that LPA-FLASH-RT produces a narrow incident energy spectrum and consequently more uniform energy deposition in the cellular layer, whereas CONV-RT exhibits a broader spectrum with greater energy heterogeneity and lower deposited energy.

2. **Dosimetry**

In this study, proton dose calibration was performed using radiochromic film (Gafchromic HD-V2). The principle relies on the dose-dependent color change of the film, which was pre-calibrated against a clinical proton accelerator using an Epson Perfection V600 scanner. RCF was placed immediately behind the cell holder, parallel to the rear Mylar window. The proton path through all absorbers (including the Kapton vacuum window, air gap, Mylar films, cell layer, and additional light-tight foils) was modelled using SRIM Monte-Carlo simulations. This yielded the ratio of dose deposition in the cell monolayer to that in the RCF. The measured RCF dose (converted to optical density from scanned images) was then corrected by this ratio to obtain the actual dose delivered to the cells.
